# Environmental DNA reveals temporal and spatial variability of invertebrate communities in arid-lands ephemeral water bodies

**DOI:** 10.1101/2024.10.31.621254

**Authors:** Brock A Hedges, Perry G Beasley-Hall, James B Dorey, Philip Weinstein, Andrew D Austin, Michelle T Guzik

## Abstract

**Context:** Throughout semi-arid and arid Australia surface freshwater is rare, and where it does occur, it is often ephemeral. This is the case for freshwater granite rock-holes that occur throughout much of southern Australia. Rock-holes support freshwater invertebrate communities, however, the ongoing threat of climate change means that this ecosystem is likely to experience hydrological disruptions. Rock-holes are also likely to be heavily impacted by invasive vertebrates. However, the ecology of this ecosystem is poorly understood despite its relative ecological significance and the extent of its associated threats.

**Aims and methods:** To provide a baseline ecological understanding of this ecosystem we documented species richness and variability at a series of rock-holes in the Gawler bioregion in South Australia using an environmental DNA approach.

**Key results:** Metabarcoding recorded invertebrates from 22 orders and 45 families. Community composition varied among rock-holes and throughout the year, with a peak in species richness in winter.

**Conclusions and implications:** These findings demonstrate the importance of these ecosystems to a range of endemic taxa. We propose establishment of monitoring programs, development of custom barcode reference libraries for the rock-hole ecosystem and future research into the likely impacts of climate change on the communities associated with them.

## Introduction

The interior of Australia is largely characterised by semi-arid and arid climates that have low annual rainfall and high rates of evaporation, resulting in a scarcity of freshwater sources (Bayly 1997). The few water bodies that are present in central Australia are often highly ephemeral in their nature and inconsistent in both their distribution and temporal persistence (Bayly 1999a, Bayly 2001). Salt lakes (May et al. 2022), ephemeral creek-lines (Reich et al. 2022), clay-pans (Gibson et al. 2018), and various water-holes and rock-holes (Timms 2013a, Timms 2014) are often among the only sources of available surface freshwater in these regions (Bayly 1999a). Freshwater granite rock-holes, present throughout much of southern Australia, are distinct from other ephemeral bodies due to their lack of connectivity and impermeable substrate (Timms 2013b, Timms 2014). These indentations in granite outcrops are formed by chemical weathering processes and provide a location for temporary storage of rainwater (Twidale and Corbin 1963, Michael et al. 2024). Periodic shifts in water levels of rock-holes are caused by the seasonality of rainfall and high evaporation rates, meaning rock-holes may alternate between being inundated to entirely dry multiple times in a single year (Timms 2014). As such, rock-holes have long been used as a source of potable freshwater for First Nations Australians (Bayly 1999a). Rock-holes are also of great value to local vertebrate species, which often migrate long distances to access freshwater (Hedges 2023). Freshwater rock-holes also act as critical refugia for a range of relictual freshwater organisms, including plants and invertebrates, that have largely disappeared from throughout the drier regions of Australia (Bayly 1997, Pinder et al. 2000, Bayly 2001). However, despite their high cultural and ecological importance, freshwater granite rock-holes are generally understudied, and little is known of their ecological roles in the wider environment (McDonald et al. 2023, Michael et al. 2024).

Invertebrate communities associated with granite rock-holes in southern Australia are consistently present despite extreme seasonal fluctuations in habitat conditions (Bayly 1997, Pinder et al. 2000, Timms 2014). These communities are dominated by crustaceans and insects (Michael et al. 2024), which persist through dry periods using adaptations likely gained during past periods of continental aridification. Adaptations include desiccation-resistant eggs, which can survive for several years without water (Bayly 1997), or free-living life stages, such as the chironomid *Paraborniella tonnoiri*, which can survive dry periods in a semi-desiccated larval state (Jones 1975). Other inhabitants of the rock-holes are less physiologically adapted to desiccation and instead rely on repeated colonisation after each rainfall event (Bayly 1997). Insects with strong flight behaviour, such as damselflies, waterbugs, and diving beetles, regularly either recolonise aquatic systems after inundation or oviposit directly into the water column (Hedges et al. 2021). These invertebrate communities are thought to provide a range of crucial ecosystem services to rock-holes through filter-feeding, which is often associated with water quality (Coughlan 1969, Atkinson et al. 2013, Buelow and Waltham 2020, Simeone et al. 2021). There is likely an association between the overall health of these aquatic systems and the diversity of resident communities of invertebrates (Kokotović et al. 2024, Suren et al. 2024).

Southern Australia is projected to experience significant drying over the next century, with all future emissions scenarios predicting a decline in rainfall and shifts in the seasonality of rainfall (IPCC 2022a). As freshwater granite rock-holes are primarily recharged by rainfall and emptied by evaporation, they will almost certainly be impacted by climate change and considerable disruptions to their historical wetting-drying cycles are expected (Bayly 1997, McDonald et al 2023). However, it is unclear how these changes in climate may impact the aforementioned invertebrate communities, as many species therein already possess adaptations to drought and desiccation.

Invertebrate communities in rock-holes throughout arid and semi-arid Australia are also threatened by invasive species. Invasive species can cause disturbance, impacting water chemistry (Doupe et al. 2010, Brim Box et al. 2016), and increasing turbidity and nitrogen concentrations (Canals 2011). Similarly, ephemeral freshwater ecosystems such as rock-holes are also particularly susceptible to invasion by algal communities (Buchberger and Stockenreiter 2018), macrophytes (Carreira et al. 2014), and macroinvertebrates (Devereaux and Mokany 2006), many of which colonise inconsistently between years. Even single-species additions or subtractions are capable of causing large and potentially detrimental changes to community composition over time (Jonsson 2006). Finally, land use and agriculture are also known to dramatically impact ephemeral water body communities (Hall et al. 2004, Dimitriou et al. 2006, Bouahim et al. 2014, Kerezsy et al. 2014, Bruno et al. 2016). All of these factors are cause for concern and are likely contributing to an overall decline in habitat security and ecosystem health of rock-holes. However, the scope of monitoring of freshwater ecosystems and their invertebrate communities in semi-arid and arid Australia is currently limited, and the freshwater rock-holes are an ecosystem that has received very little study (Michael et al. 2024).

The Government of South Australia has identified sampling of the macroinvertebrate communities associated with the rock-holes in the Gawler bioregion as a priority action to assess their ecological value (White 2009). However, the remoteness of semi-arid and arid Australia and a lack of infrastructure, means that targeted biomonitoring programs are difficult to undertake (McDonald et al. 2023). Previous surveys have used traditional ecological techniques, involving direct specimen collection and morphological identification, to estimate rock-hole community diversity and abundance (Timms 2014, Pinder et al. 2000). Since taxonomic expertise is increasingly rare, particularly for invertebrates (Yeates et al. 2003, Austin et al. 2004, Engel et al. 2021), emerging technologies have gained considerable interest as viable options for assessing ecosystem health. Environmental DNA (eDNA) is one such tool that involves the collection of environmental samples, such as soil or water, and bulk amplification of shed genetic material from organisms that might, or might not, be physically present at the time of sampling. This technique is relatively non-destructive and non-invasive as communities can be sampled and assessed with minimal disturbance and removal of material (White et al. 2020). Environmental DNA has recently been used to improve detection, monitoring, management, and conservation of endangered freshwater taxa (Rodgers et al. 2020), as well as to characterise community composition (Holman et al. 2019). It has been used to detect species present at low densities (Johnsen et al. 2020), non-indigenous and invasive invertebrate species (Holman et al. 2019), and ecosystem stressors (Fan et al. 2020). These methods rely on barcode reference libraries (BRLs), which assign taxonomy to otherwise anonymous DNA sequences and therefore provide biological meaning to eDNA metabarcoding reads and operational taxonomic units (Rimet et al. 2021, Michael et al. 2024). Metabarcoding is one common method of characterising eDNA and involves the amplification of a single genetic marker, or “barcode”, using high-throughput sequencing technology. When BRLs are robust and contain extensive information on target taxa, eDNA metabarcoding can identify species more accurately than traditional morphological approaches (Galimberti et al. 2021). Environmental DNA therefore represents a rapid and robust method for identifying species for which taxonomic expertise may be lacking, such as the invertebrate communities found in Australian freshwater rock-holes. Most Australian rock-hole research that has used eDNA methods have focussed on vertebrates (McDonald et al. 2023, Mousavi-Derazmahalleh et al. 2023, Hedges et al. 2024a) with only one study focussed on invertebrate communities, and this in a temperate climatic zone (Michael et al. 2024). To date, the invertebrate communities associated with the granite rock-hole ecosystems of semi-arid and arid Australia have only been surveyed using traditional methods (Bayly 1997, Pinder et al. 2000, Timms 2014).

To address these knowledge gaps, we tested the application of eDNA metabarcoding for detecting ephemeral freshwater invertebrates. The specific aims of this research were to a) test eDNA metabarcoding as a tool for detecting invertebrates in granite rock-holes of South Australia; b) characterise the composition of invertebrate communities in these ecosystems; c) determine whether these communities varied spatially and temporally; and d) provide recommendations regarding community conservation and management. It is predicted that eDNA metabarcoding will allow detection of a suite of invertebrate taxa, including crustaceans and insects. It is further predicted that the communities recovered with eDNA metabarcoding will vary spatially and temporally.

## Materials and methods

### Site description

Hiltaba Nature Reserve (HNR) is a large ex-pastoral property that borders the Gawler Ranges National Park to the north of the Eyre Peninsula in South Australia (Figure 1). The Reserve is approximately 78,000 ha and has been managed for conservation outcomes by Nature Foundation since its acquisition by the organisation in 2012. The Reserve is primarily composed of a series of rolling granite hills interspersed with woodland and grassland and contains unique habitats used by local plants and animals known to be in decline elsewhere (Nature Foundation 2023). The Nature Foundation has enacted a series of conservation programs that aim to improve biodiversity management at HNR.

**Figure 1.**
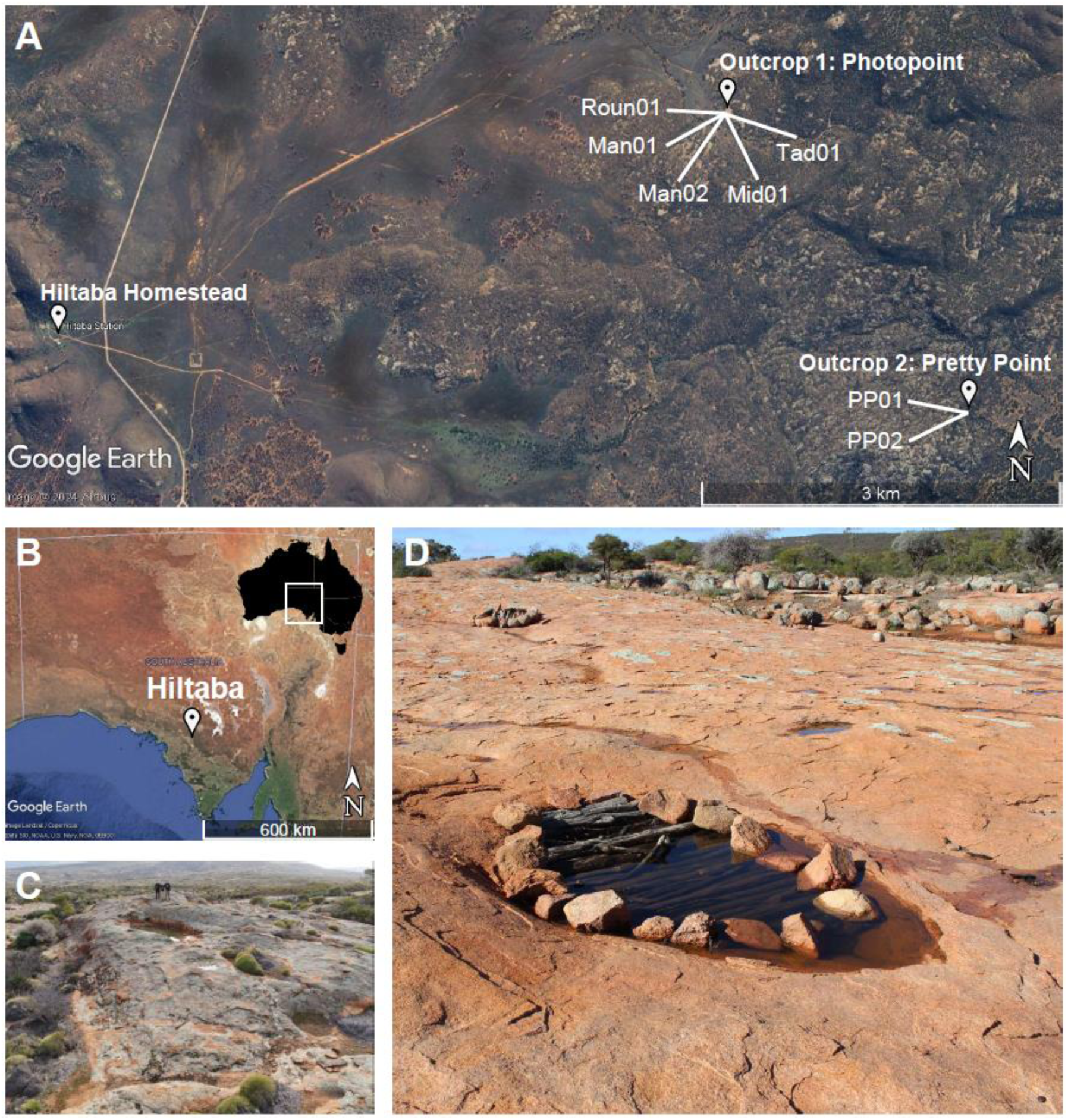
Study location of the present study. A) locations of the seven rock-holes (Table 1) sampled for freshwater eDNA within Hiltaba Nature Reserve (HNR); B) The location of HNR in South Australia, with the location of South Australia within the Australian continent depicted inset; C) a rain-filled granite rock-hole at the ‘Pretty Point’ outcrop at HNR; D) a rain-filled granite rock-hole on the ‘Photopoint’ outcrop at HNR, diameter of the rock-hole at widest point = 2.2 m.

**Table 1.** Summary of the rock-holes sampled for freshwater during February, July, and October of 2020 (green checkmark). Rock-holes not sampled at certain time points (red cross) were either empty or had only a small volume of water present. Rock-hole type assignments have been adapted from Timms (2013b). Outcrop 1 refers to the Photopoint outcrop, and outcrop 2 refers to the Pretty Point outcrop.

| Rock-hole features |  |  |  |  |  | Sample period |  |  |
| --- | --- | --- | --- | --- | --- | --- | --- | --- |
| ID | Latitude | Longitude | Elevation (m) | Type | Outcrop | Feb | July | Oct |
| Man01 | -32.13941 | 135.13461 | 216 | Pit | 1 | ✓ | ✓ | ✓ |
| Man02 | -32.13943 | 135.13478 | 216 | Pit | 1 | ✓ | ✓ | ✗ |
| Tad01 | -32.1395 | 135.1348 | 216 | Pit | 1 | ✓ | ✓ | ✓ |
| Roun01 | -32.13931 | 135.13492 | 217 | Pan | 1 | ✓ | ✓ | ✓ |
| Mid01 | -32.13941 | 135.13461 | 217 | Pit | 1 | ✓ | ✓ | ✓ |
| PP01 | -32.165521 | 135.1488 | 312 | Pan | 2 | ✗ | ✓ | ✓ |
| PP02 | -32.165521 | 135.1488 | 310 | Pan | 2 | ✗ | ✓ | ✗ |

The granite hills present throughout much of HNR are entirely exposed in many locations, resulting in granite outcrops of varying slope and morphology. These outcrops are generally flat with slopes of <20° and often display depressions which, following rain, provide an impermeable location for storage of water and sediments. These depressions are considered to be rock-holes, consistent with those discussed above, and support a suite of vertebrates (Hedges 2023), aquatic invertebrates and plants.

### eDNA sampling

Freshwater eDNA sampling was undertaken in 2020 at seven rock-holes in HNR on February 11–12 (summer), July 9–11 (winter), and October 2–3 (spring) (Table 1). Sampling of seven rock-holes was undertaken across two rocky outcrops approximately 3 km apart (Figure 1).

Four of these rock-holes were characterised as pit rock-holes and three were pan rock-holes following the classification scheme by Timms (2013b), which defines pans as “of diverse shape in plan, shallow, flat-floored and seasonally filled with water”. This is in contrast to pits which are described as “typically subcircular in plan, have a depth to diameter ratio exceeding 0.2, and contain water for longer periods”. During the summer time point (February), only rock-holes at outcrop 1 were sampled due to safety considerations associated with summer sampling of rock-holes at outcrop 2 (Table 1). During the spring time point (October), rock-holes ‘Man02’ and ‘PP02’ were not sampled due an insufficient volume of standing water.

Five 1 L replicates of water were collected from each rock-hole. Five replicates have previously been shown to be sufficient for detection within a range of freshwater systems (Shaw et al. 2016). One blank sample was taken at each rock-hole and filled using 1 L of store-bought bottled water. Replicates were collected using one-litre, wide mouth NALGENE^TM^ bottles that were sterilised with 20% bleach and dried prior to use. Samples were collected whilst wearing disposable latex gloves, stored in and transported in clean plastic tubs to minimise contamination. Sampling bottles were cleaned first with bleach solution, and then ethanol before reuse. Negative field controls (prepared in the field but using RO water) and equipment controls (prepared in the filtering room using RO water) were used to test for contamination.

Replicates and blanks were filtered on-site through membranes with 0.44 µm pores using a JAVAC vacuum pump (model CC-45) in February 2020. The pump was connected to a series of conical flasks, the first filled with silica beads. The second conical flask was attached to a magnetic filter funnel. A Pall Sentino microbiology pump was used in July and October 2020. Membranes were transported on ice and stored at -20°C prior to DNA extraction. A pore size of 0.44 µm was selected due to the high concentration of suspended solids present in rock-hole freshwater samples. Pump equipment was wiped down with bleach solution and then ethanol between filtering for each rock-hole.

### eDNA laboratory methods

We extracted DNA from half of each filter paper using a modified Qiagen DNeasy blood and tissue kit protocol (Qiagen, Germany) and an automated QIAcube extraction platform (Qiagen). Where more than one filter paper was used for a sample, equal portions of each paper were taken to a total of half of a filter paper. All extractions were undertaken in a dedicated PCR-free laboratory, and extraction controls were processed alongside samples. Extractions were eluted in a final volume of 100 µL AE buffer.

To determine the required dilution for optimal amplification, polymerase chain reactions (PCRs) were performed in duplicate on each extraction by adding DNA template directly to the PCR master mix (neat), then performing a serial dilution (1:10). We used proprietary cytochrome oxidase subunit I (*COI*) insect-mollusc primers and 16S crustacean primers. The PCRs were performed at a final volume of 25 µL where each reaction comprised of: 1× PCR Gold Buffer (Applied Biosystems), 0.25 mM dNTP mix (Astral Scientific, Australia), 2 mM MgCl_2_ (Applied Biosystems), 1 U AmpliTaq Gold DNA polymerase (Applied Biosystems), 0.4 mg/mL bovine serum albumin (Fisher Biotec), 0.4 µM forward and reverse primers, 0.6 μL of a 1:10,000 solution of SYBR Green dye (Life Technologies), and 2 µL template DNA. A series of PCRs were performed on StepOne Plus instruments (Applied Biosystems) with the following cycling conditions: 95°C for 5 min, followed by 50 cycles of: denaturation at 95°C for 30 sec, annealing at 49°C for 30 sec, extension at 72°C for 45 sec, then a melt-curve analysis of: 95°C for 15 sec, 60°C for 1 min, 95°C for 15 sec, finishing with a final extension stage at 72°C for 10 min.

After selection of the optimal dilution (neat or 1:10), PCRs were repeated in duplicate as described above using unique, single-use combinations of 8 base pair (bp) multiplex identifier-tagged (MID-tag) primers as described in Koziol et al. (2019) and van der Heyde et al. (2020). Master mixes were prepared using a QIAgility instrument (Qiagen) in an ultra-clean lab facility, with negative and positive PCR controls included on each plate. A sequencing library was constructed by combining samples into mini-pools based on the PCR amplification results from each sample. The mini-pools were analysed using a QIAxcel (Qiagen) and combined in roughly equimolar concentrations to form libraries. Libraries were size selected (250–600 bp cut-off) using a Pippin Prep instrument (Sage Sciences) with 2% dye-free cassettes, cleaned using a QIAquick PCR purification kit, quantified on a Qubit (Thermo Fisher), and diluted to 2 nM. Libraries were sequenced on an Illumina MiSeq instrument using a 500-cycle V2 kit with custom sequencing primers.

### Bioinformatics and statistical analyses

Metabarcoding data were processed using QIIME2 v.2021.11 (Bolyen et al. 2019). Raw sequences were demultiplexed, trimmed by quality scores (a subset of samples was visualised in QIIME2 View and a consensus trim length selected by eye), filtered to remove chimeric sequences, and denoised using DADA2, which accounts for amplification errors (Callahan et al. 2016). Sequences were then tabulated to construct ZOTUs (zero-radius operational taxonomic units), which have a clustering threshold of 100%, as a proxy for species and to derive representative sequences for each ZOTU. Representative sequences were aligned using MAFFT (Katoh et al. 2002) and a midpoint-rooted phylogeny was generated using FastTree 2 for downstream calculation of diversity metrics (Price et al. 2010). A custom database was constructed from all invertebrate *16S* rRNA and *COI* sequences available as of June 9, 2022 via the online GenBank repository (National Center for Biotechnology Information) using the following search strategy: ((("16S ribosomal RNA" OR "16S rRNA" OR mitochondrion OR “CO1” OR “COI” OR “cytochrome oxidase subunit I” OR “cytochrome oxidase subunit 1”) AND (Mollusca[Organism] OR Arthropoda[Organism]) NOT Vertebrata[Organism] OR Archaea[Organism] OR Fungi[Organism]). We then used BLASTN (Camacho et al. 2009) with default settings applied, to query our representative sequences against the custom database with the maximum number of target sequences set to 1 per query. Representative sequences with non-invertebrate top hits, or those without a top hit that had a corresponding species-level assignment, were excluded from further analysis. The filtered dataset was then subjected to alpha diversity analyses using metrics of evenness (Pielou 1966), Faith’s phylogenetic diversity (Faith 1992), and Shannon’s diversity index (Shannon and Weaver 1949). As we were interested in examining difference in community composition on spatial and temporal levels, we also conducted beta diversity analyses using Bray-Curtis (Bray and Curtis 1957), Jaccard (Jaccard 1901), and weighted and unweighted Unifrac distances (Lozupone et al. 2011) against rock-hole location and sample collection date. Principle coordinate analysis (PCoA) plots of these metrics were exported to R version 4.2.2 (RCoreTeam 2013) using QIIME2R v0.99.6 for further visualisation (Bisanz 2018). Stacked bar plots, violin plots, and PCoA plots were generated using *ggplot2* v3.4.0 (Wickham 2016). Rarefaction curves were generated using *iNEXT* v3.0.0 (Hsieh et al. 2022). UpSet plots were generated using *UpSetR* v1.4.0 (Conway et al. 2017).

## Results

### Environmental DNA data

We successfully recovered invertebrate eDNA from 40 of 91 samples using the *COI* insect-mollusc assay (see above). In total, we generated 5,379,229 sequences with 116,939 mean sequences per sample. A total of 179 ZOTUs were detected using BLASTN hits from GenBank. Of these, seven were present only in blanks and positive controls and a further 113 were considered unlikely to belong to freshwater invertebrate taxa after cross-referencing with online repositories (Atlas of Living Australia, World Register of Marine Species), and making a case-by-case decision regarding their likelihood of being aquatic. Excluded taxa, which often belonged to groups of flying insects like Hymenoptera and Hemiptera, were excluded from downstream analysis. The remaining 59 ZOTUs belonged to 41 families within 20 orders (Figure 2A).

**Figure 2.**
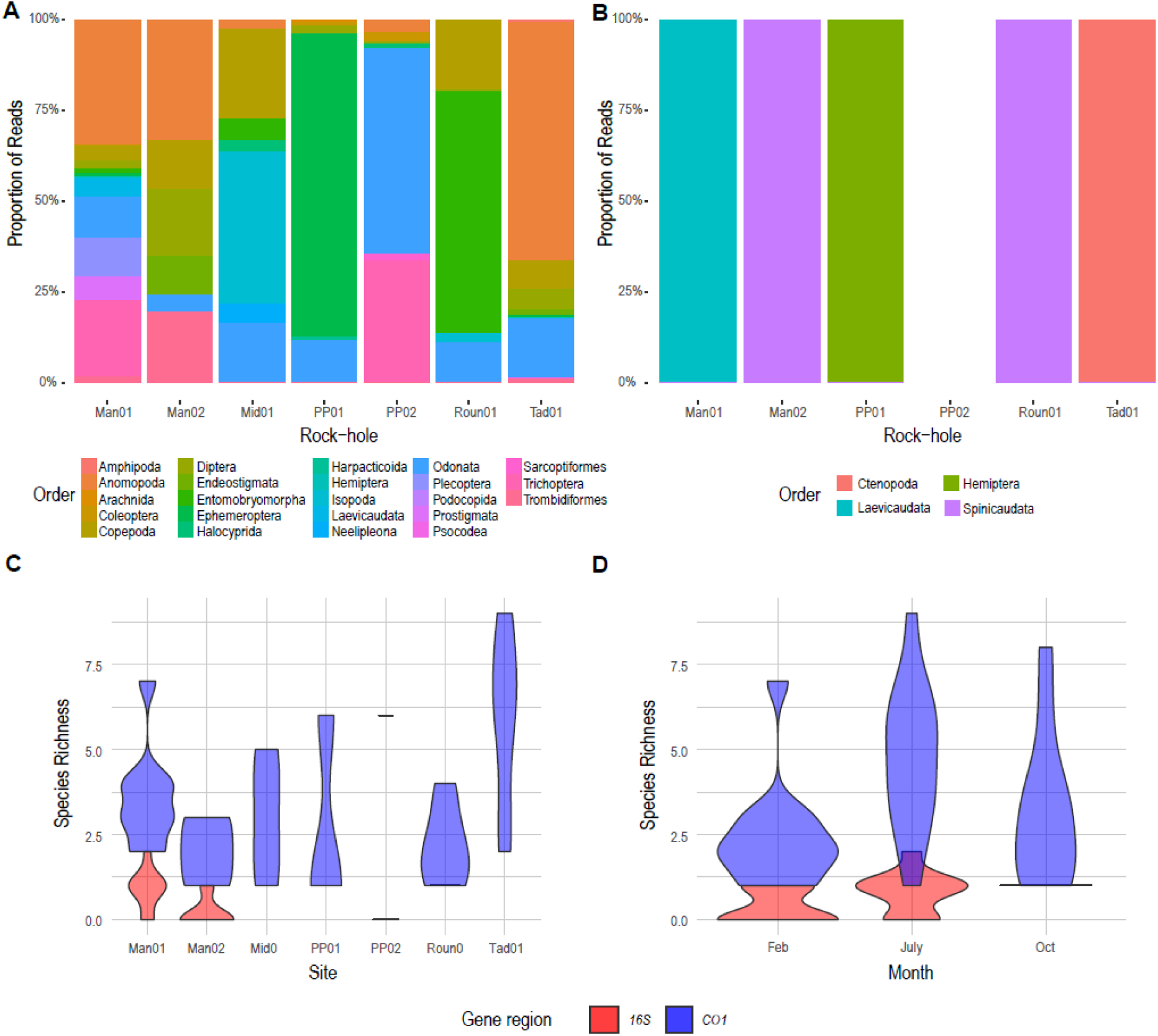
A) Stacked bar plots depicting the proportion of sequences assigned to family for each rock-hole for COI; B) stacked bar plots depicting the proportion of sequences assigned to family for each rock-hole for 16S rRNA; C) violin plots showing variation in species richness among rock-holes for COI (blue) and 16S rRNA (red); and D) violin plots showing variation in species richness among sample collection periods (month at which eDNA samples were collected) for COI (blue) and 16S rRNA (red). The full dataset is presented in appendix Tables 1 and 2, and stacked bar charts for the distribution of reads by family can be seen in Figure S1.

In contrast, the *16S* rRNA crustacean assay recovered invertebrate eDNA from 41 of 91 samples, yielding 5,758,543 sequences with 122,522 mean sequences per sample. A total of 35 ZOTUs were detected using BLASTN hits from the GenBank-derived BRL. Of these, seven were present only in blanks and positive controls, and 16 were considered unlikely to belong to freshwater invertebrate taxa and were excluded from downstream analysis. The remaining 13 ZOTUs belonged to 4 families each within their own order (Figure 2B).

Between both datasets, invertebrates were recorded from 45 families within 22 orders with an average BLASTN pairwise identity match of 89%.

Through comparison with the species lists presented by Timms (2014) we found 30 families and 14 orders that had not previously been recovered from South Australian freshwater granite rock-holes were detected using eDNA metabarcoding (Hedges et al 2024b, via figshare). Notable new order and suborder records included stoneflies (Plecoptera), harpacticoid copepods (Harpacticoida) and various mites (Sarcoptiformes, Trombidiformes, Prostigmata, and Endeostigmata). Overall ZOTU species richness for the sequences with top hits to GenBank records matches did not vary noticeably among sites (Figure 2C) and instead more considerably by sampling time, with winter/spring (July/October) collections displaying greater ZOTU richness than the summer collection period (Figure 2D).

Rarefaction could only be performed for five timepoints for the *COI* dataset and none of the replicates for *16S* rRNA due to an insufficient number of data-rich replicates. Rarefaction of species richness indicated that 60–75% of the diversity was sampled and that further sampling would likely improve capture to over 75% of invertebrate diversity (Figure S2). Whilst species richness was relatively consistent across sites, ZOTUs were highly variable, with the majority of taxa (37 ZOTUs for COI and 3 ZOTUs for 16S rRNA) occurring only in a single rock-hole (Figure S3).

Principle coordinate analysis showed that replicates tended to cluster closest to other replicates from the same rock-hole (Figure 3). Collection month had a slight effect, although this was most pronounced for *16S* samples before the taxonomic filter was applied. In the *COI* dataset and for *16S* rRNA after the taxonomic filter had been applied, there was a degree of overlap in multivariate space across the collection months (Figure 3).

**Figure 3.**
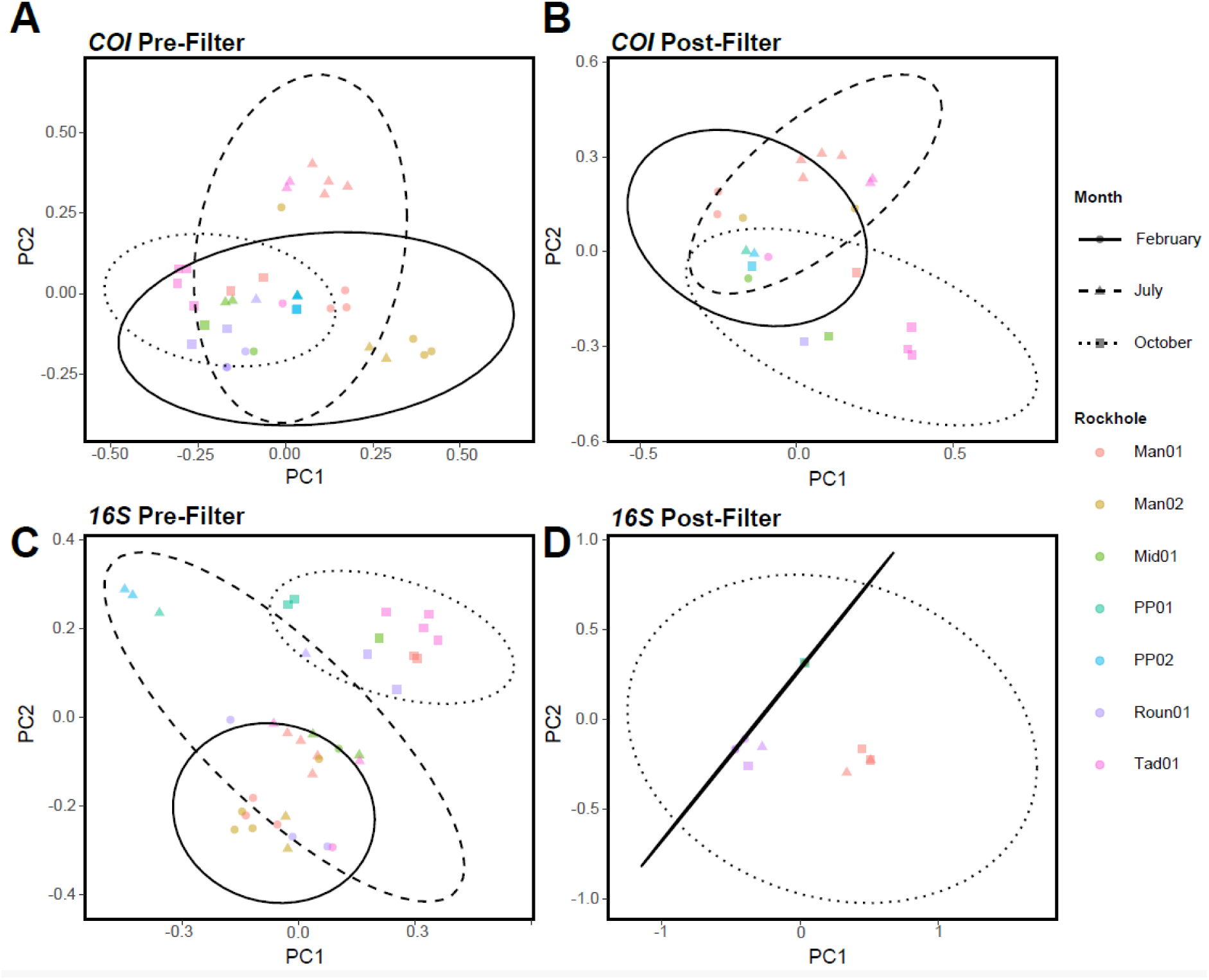
PCoA plots showing maximum dissimilarity for pre-filter (all sequences included) and post-filter (non-invertebrate sequences excluded) eDNA samples based on Jaccard distances. Principle component (PC) 1 and PC2 represent maximum dissimilarity between samples in order of magnitude. Colours represent individual rock-holes and shapes represent the month during which eDNA sampling was undertaken. Ellipses have been applied to show the 95% confidence interval (CI) for similarity between sample collection months.

## Discussion

Here we present the first molecular ecological assessment of the invertebrate communities of semi-arid and arid rock-hole ecosystems using eDNA metabarcoding approaches. This is the second such assessment for Australia rock-holes more broadly (Michael et al. 2024).

Freshwater invertebrate species from 22 orders and 45 families in seven freshwater granite rock-holes at HNR in southern Australia were detected. Traditional ecological methods have been employed to survey invertebrate communities in rock-holes throughout southern Australia (Bayly 1997, Pinder et al. 2000, Timms 2014, Hedges et al. 2021), but the present study is the first to employ eDNA as a more rapid tool for assessing community composition. Identification of lineages within rock-holes to family was generally possible, but barcode reference libraries generated from GenBank records were insufficient for species-level identification due to a lack of barcodes deposited from closely-related taxa. Invertebrate communities did not vary dramatically across HNR, although they did shift throughout the year by season, with species richness peaking in winter. Below, we discuss the suitability of environmental DNA as a tool for biomonitoring the freshwater granite rock-hole ecosystem and present a series of conservation recommendations. Past studies on granite rock-holes elsewhere in southern Australia have discovered high rates of rarity and short-range endemism among invertebrates (Bayly 1997, Pinder et al. 2000).

The rock-holes at HNR are likely to be amongst the most pristine in the region as other areas of the Gawler bioregion are still actively stocked with sheep, with only HNR and the neighbouring Gawler Ranges National Park entirely destocked (Nature Foundation 2023). It is therefore likely that rock-holes elsewhere are far more highly impacted than at the study site examined here. Understanding changes in invertebrate community structure of rock-holes, in particular indicator species which are typically susceptible to environmental change, such as chironomids (Czechowski et al. 2020, Jones 1975, Pinder et al. 2000), are critical to estimating the ecological significance of rock-holes more generally (Jenkin et al. 2011). Of the 22 orders and 45 families we identified from sequences available on the online GenBank repository (National Center for Biotechnology Information), a number of common freshwater bioindicator groups (Firmiano et al. 2017, Czechowski et al. 2020, Schröder et al. 2020, Vilenica et al. 2021) were detected including chironomids (Diptera: Chironomidae), mayflies (Ephemeroptera: Baetidae), and dragonflies (Odonata) (see Figure 2, S1, Table S1 (Hedges et al 2024b, via figshare)). These groups occurred frequently across samples, but feature counts (the number of sequences that mapped to a species) varied dramatically. Interestingly, the most dominant organismal groups in terms of relative read abundance were Anomopoda (water fleas) for *COI* and Spinicaudata (clam shrimps) for *16S* rRNA. Fourteen orders and 30 families had not previously been detected in South Australian freshwater granite rock-holes (Timms 2014, see Table S3 for the complete comparison (Hedges et al 2024b, via figshare). Notable new order and suborder records included stoneflies (Plecoptera), Harpacticoid copepods (Harpacticoida), and various mites (Sarcoptiformes, Trombidiformes, Prostigmata, Endeostigmata). These records contribute new information to our understanding of the composition of rock-hole invertebrate communities in semi-arid and arid Australia. Several of these groups are known to occur in freshwater rock-hole ecosystems of Western Australia (Pinder et al. 2000, Bayly 1997), a system thought to have been isolated from the South Australian rock-holes by the biogeographic barrier of the Nullarbor Plain (Timms 2014).

Validation of these records through targeted field surveys would allow for further identification and incorporation into barcode reference libraries to facilitate future metabarcoding work.

Rock-holes, such as those in granite discussed here, have been previously proposed as ecosystems of significant evolutionary value, both in the short-term (as refuges) and evolutionary timescales (as refugia). *Refuges* are locations that allow biota to persist despite short-term conditions of disturbance, such as drought or flooding, whilst *refugia* are locations that allow biota to persist despite environmental change occurring across longer evolutionary time periods (Davis et al. 2013). Davis et al. (2013) further qualified the distinction between refuges and refugia by proposing that temporary and ephemeral water bodies are likely to act as *refuges* for only highly mobile taxa—such as dragonflies and water beetles—but may serve as *refugia* for less mobile taxa, such as less dispersive freshwater crustaceans. We detected invertebrate lineages belonging to both of these categories in the present study. The rock-holes of the Gawler Bioregion may therefore represent locations from which individuals present in source populations may disperse to replenish the egg (and seed) banks of nearby water bodies (Sheldon et al. 2010, Davis et al. 2013). Our characterisation of these communities using eDNA metabarcoding builds upon the work of Davis et al. (2013) by improving our understanding of both the composition of these communities and their variability at spatial and temporal scales.

### Insights into spatial and temporal community variability

The results described here provide insights into invertebrate community dynamics of Australian freshwater rock-holes at spatial and temporal scales. Overall species richness was not observed to vary dramatically among rock-holes (Figure S3), but community composition varied significantly (Figures 2A, 2B, 3). The rock-holes also showed a degree of spatial variability regarding community composition (Figure 3). Interestingly, we observed no effect of outcrop location, with rock-holes from both outcrops nested among one another. This finding is consistent with that of Pinder et al. (2000) and Timms (2014) for similar rock-holes elsewhere in Australia, where only slight clustering of rock-holes by outcrop was observed based on invertebrate assemblages. This may suggest that efforts to conserve rock-hole biodiversity need not consider small geographical distances between rock-holes, and should instead focus at landscape-wide efforts. Although we only surveyed two outcrops on one property in the present study, rock-holes are found throughout the Gawler Bioregion (Jenkin et al. 2011). Sampling of additional rock-holes spanning the entirety of the Gawler Bioregion may improve confidence in assessments of spatial variability.

Rock-hole invertebrate communities varied temporally in their composition based on our eDNA metabarcoding data, with species richness peaking in winter across all samples (July) (Figure 2C, 2D). Community composition varied by sampling month, with some overlap in communities occurring in both the pre- and post-filter dataset (Figure 3). Whilst it is likely that sampling throughout a single year is insufficient to accurately capture temporal variation in these communities, our results support previous research on temporal variability in rock- hole and other ephemeral freshwater communities (Bayly 2001, Timms 2014) indicating substantial variability of the presence of particular taxa within and between years. In particular, Timms (2014) showed that five visits over two to three years were required for species accumulation curves to reach a plateau, which would suggest that the majority of detectable species had been recorded. Similarly, Bayly (2001) observed gradual change in species composition over time within a single freshwater rock-hole. Given these observations, we suggest successive sampling over consecutive years would improve confidence that the entire macroinvertebrate community has been adequately sampled. This inference is supported by rarefaction, suggesting that additional field seasons are likely detect additional taxa not recorded here (Figure S2).

### Environmental DNA as a biomonitoring tool

Environmental DNA metabarcoding generally performs well when compared to more traditional methods for assessing freshwater invertebrate communities (Robson et al. 2016, van der Heyde et al. 2023, Keck et al. 2022, Villacorta-Rath et al. 2022, Davis et al. 2023). McDonald (2023) used freshwater eDNA deposited by vertebrates to assess rates of visitation to freshwater granite rock-holes. Here, we have shown the common macroinvertebrate taxon groups present at HNR include Anomopoda (water fleas), Diptera (flies), Hemiptera (true bugs), Odonata (damselflies and dragonflies), and Sarcoptiformes (mites) and are consistent with survey results and morphological identifications performed by Timms (2014).

Traditional survey techniques often rely on removal of material from the system, including in many cases live organisms (Timms 2014, Hedges et al. 2021). For systems of relatively small volume that are slowly recharged by water, such removal can be disruptive and destructive.

The advantage of sampling freshwater eDNA is that only a small volume of water is required from the system (McDonald et al. 2023) and without destructive sampling of live organisms. Additionally, water sampling/collection is easy and can be undertaken by field operatives (e.g., landholders, conservation workers, and volunteers) without substantial scientific training (Prie et al. 2021, 2023). However, water availability at these locations can be limited at certain times of the year and may impact capacity to apply freshwater eDNA collection methods at times. As such, we suggest that eDNA metabarcoding is a promising method for detecting invertebrates in freshwater bodies such as rock-holes.

In our study, incomplete barcode reference libraries substantially limited the extent to which metabarcoding could determine robust species names for ZOTUs. This observation is not uncommon for freshwater ecosystems (Michael et al. 2024) and even for comparatively well-studied freshwater invertebrate taxa, such as Odonata (dragonflies and damselflies), reference datasets are often lacking (Galimberti et al. 2021), and only allow identification to higher taxonomic levels (Mulero et al. 2021). As a result, inferences made regarding the functional roles of species are limited by low confidence in species identities (Zhang et al. 2023). More robust barcode reference libraries would improve this confidence and allow for assessments of functional groups (Rimet et al. 2021). Comprehensive BRLs improve confidence in species assignments and in eDNA metabarcoding studies generally (Ekrem et al. 2007, Weigand et al. 2019, Saccò et al. 2022). As such, we recommend future research to develop BRLs for the granite rock-hole ecosystem.

## Conclusions

### Future work

Applications of eDNA metabarcoding are currently limited in the scope of ecologically informative data that can be gathered, due partly to the incompleteness of BRLs as discussed above. However, other limitations impact the broad application if eDNA metabarcoding methods such as a disconnect between eDNA sequence abundance and both organismal abundance and biomass, which limits ecological inference regarding these factors (Johnsen et al. 2020). Studies requiring these data often need to pair eDNA collections with more traditional methods such as isotopic and radiocarbon analyses which allow inference of diet regimes, food webs, and ecosystem function (Saccò et al. 2019) and traditional sampling surveys that count abundance directly. The present study has demonstrated the validity of eDNA as a broader tool, suited to assessing the various communities associated with freshwater ecosystems. In the future, robust assessment of specific taxa and more targeted freshwater eDNA approaches making use of taxon-specific assays would provide more detailed and monitoring-relevant information. We also recommend that such surveys be used as an opportunity to collect specimens to assemble more robust and ecosystem-relevant BRLs (Rimet et al. 2021).

In some cases, eDNA metabarcoding allows for assessment of the impact of environmental factors such as dissolved oxygen and heavy metals on invertebrate communities (Wang et al. 2023), as well as in making inferences regarding potability of drinking water (Shim et al. 2023). As the rock-holes are a source of drinking water to local vertebrate communities (McDonald et al. 2023, Hedges 2023), adapting techniques could allow for assessment of resource quality in conservation programs.

Our findings demonstrate the suitability of eDNA metabarcoding as a method for determining community composition of freshwater rock-holes, in this case in the Gawler Bioregion. We were able to identify key taxonomic groups living in these communities, many of which represented new records for this system outside of Western Australia. We gauged the variability at spatial and temporal scales of these communities, which saw a peak in species richness during winter. Invertebrate communities in rock-holes on different outcrops were comparable to one another, which suggests that conservation management directed at alleviating the impacts of climate change and invasive species may be best applied at a landscape scale, rather than through targeting individual rock-holes. We recommend that eDNA metabarcoding be used as a method for monitoring these communities and that future research be undertaken to improve barcode reference libraries to facilitate such work.

## Data Availability Statement

All data used in the preparation of this manuscript have been made available online through the appropriate data repositories. Environmental DNA metabarcoding data and summary tables can be accessed through figshare at: https://figshare.com/projects/Environmental_DNA_reveals_temporal_and_spatial_variability_of_invertebrate_communities_in_arid-lands_ephemeral_water_bodies/226008

## Acknowledgements

We would like to thank The University of Adelaide, Nature Foundation and the Ecological Society of Australia for financial support. Thanks to colleagues (Oliver Gore, Dr Jenna Draper, Alana McClelland, Dr Adam Toomes, Dr Nicole Foster, Johanna Kuhne, Charlie Wilson, Taylah Wallach and Lachlan Perry) for assisting with field data collection. Thanks to the Nature Foundation rotational management team for assistance and support whilst on field work, as well as Alex Nankivell, for his eagerness to share invaluable experience in the field.

## Supplementary Material

**Figure S1.**
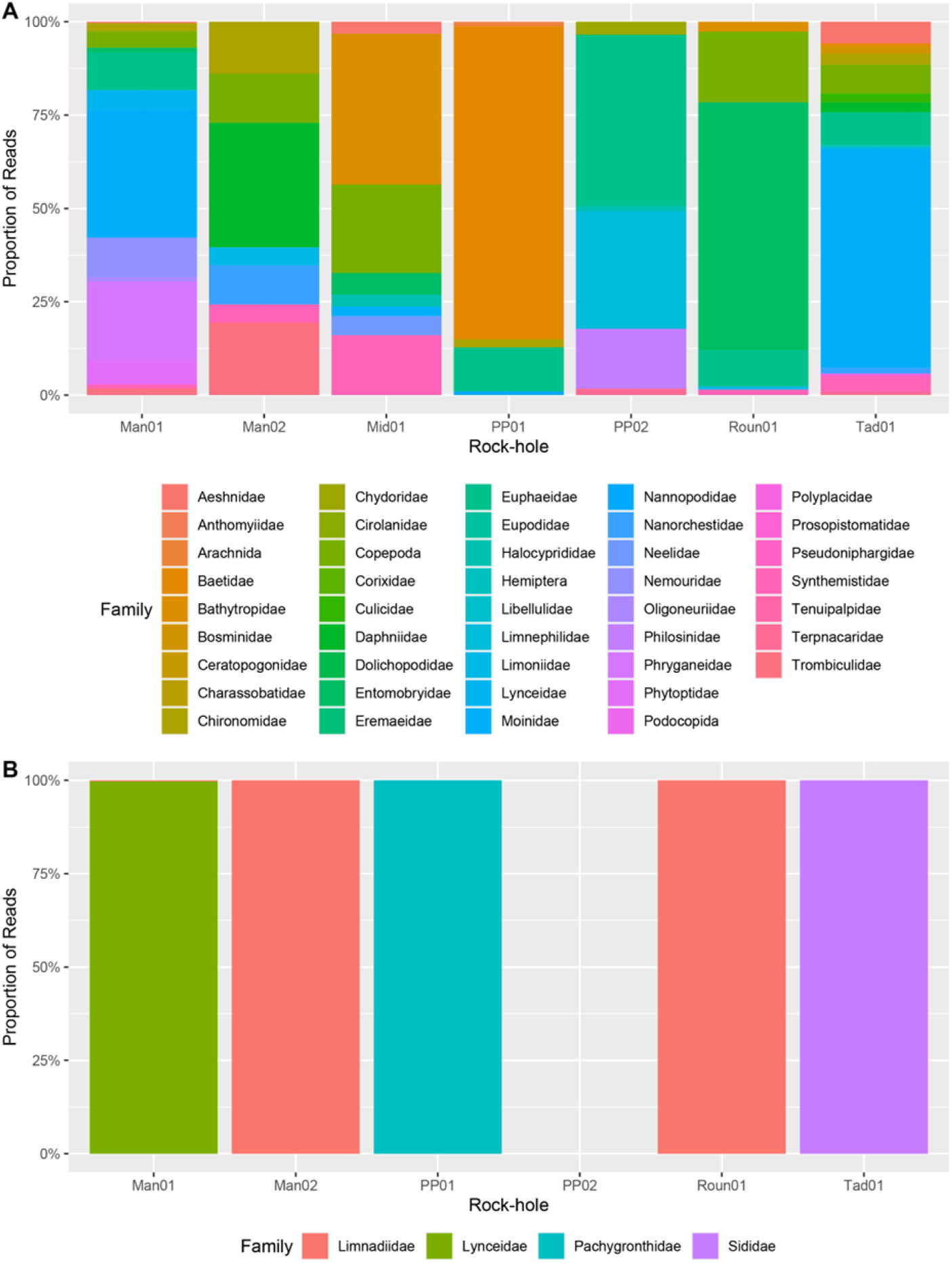
Stacked bar plots depicting the proportion of sequences assigned to family for each rock-hole for A) *COI* and B) *16S* rRNA. The full dataset is presented in Table S1 (Hedges et al 2024b, via figshare), and stacked bar charts for the distribution of reads by order can be seen in Figure 4.2. Rock-hole Mid01 has been excluded from data presented in B due to sample failure.

**Figure S2.**
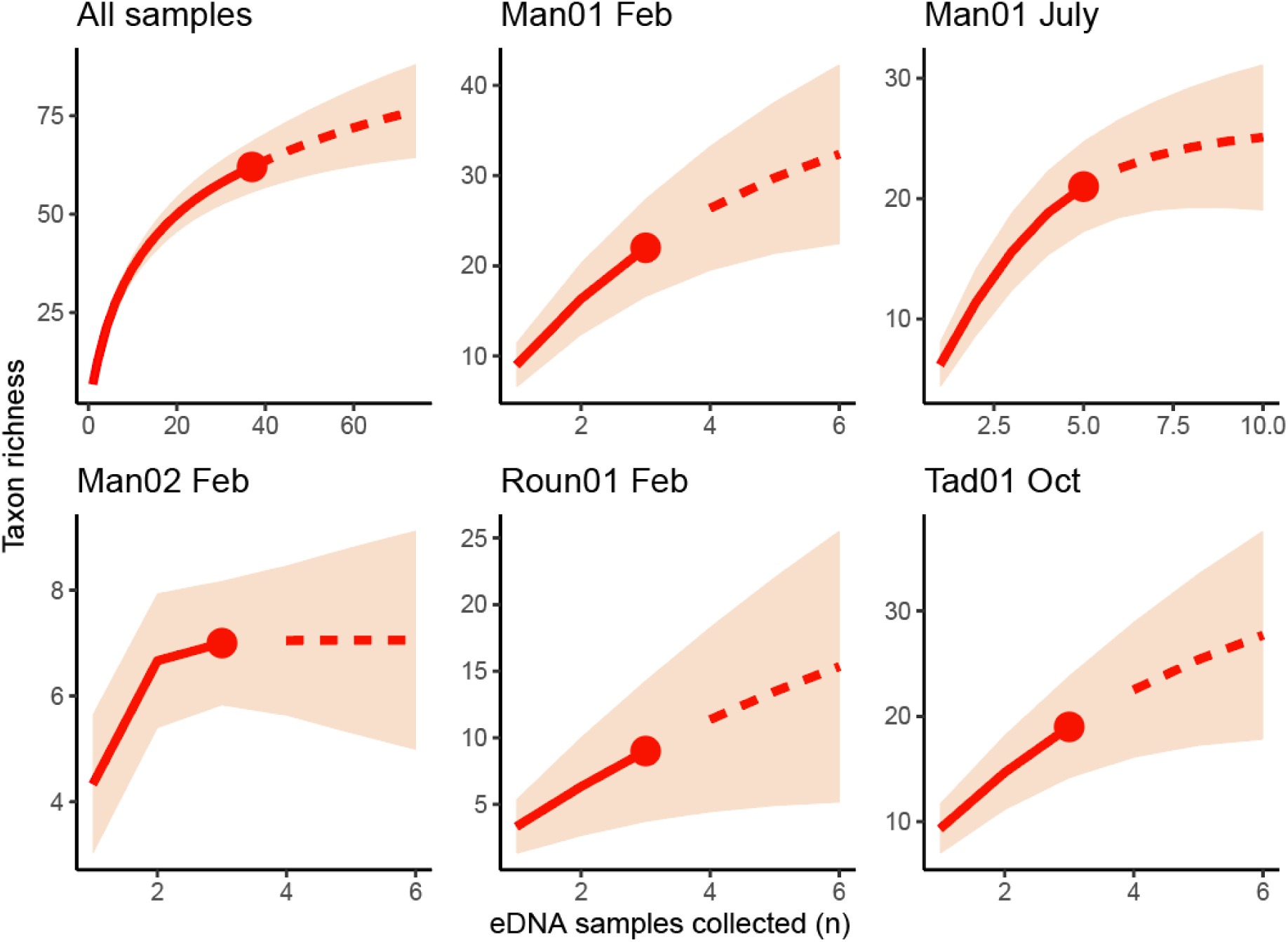
Rarefaction plots showing the impact of replication of freshwater eDNA samples for the entire dataset (top left) and at different rock-holes.

**Figure S3.**
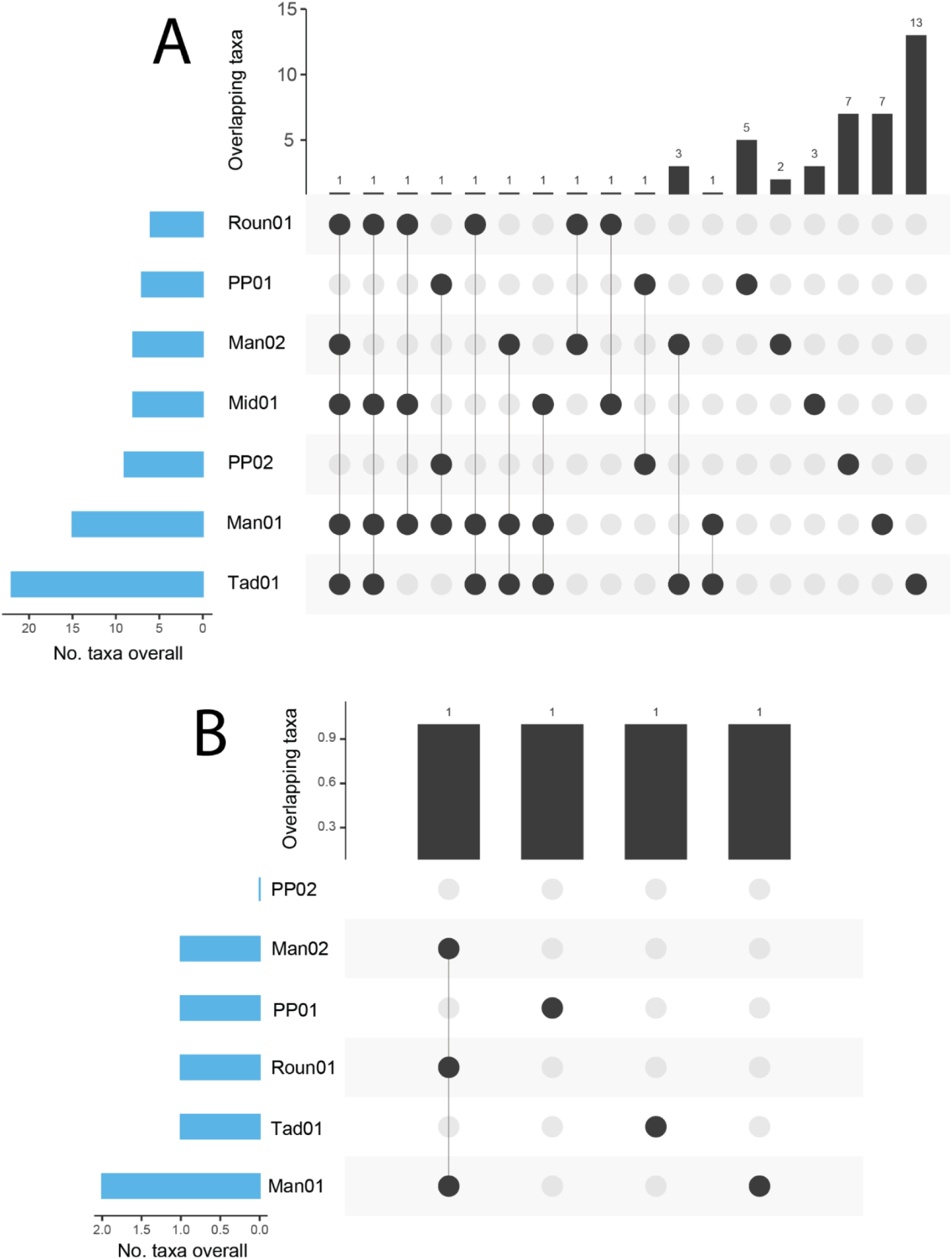
UpSet plots showing the number of ZOTUs shared among rock-holes for A) *CO1* data; and B) *16S* rRNA data. Blue bars represent the total number of taxa within a single site. Black bars represent taxa that are shared between each site with black circles indicating which sites these taxa are shared with. UpSet plots showing the data without replicates collapsed can be seen in Figures S4 and S5.

**Figure S4.**
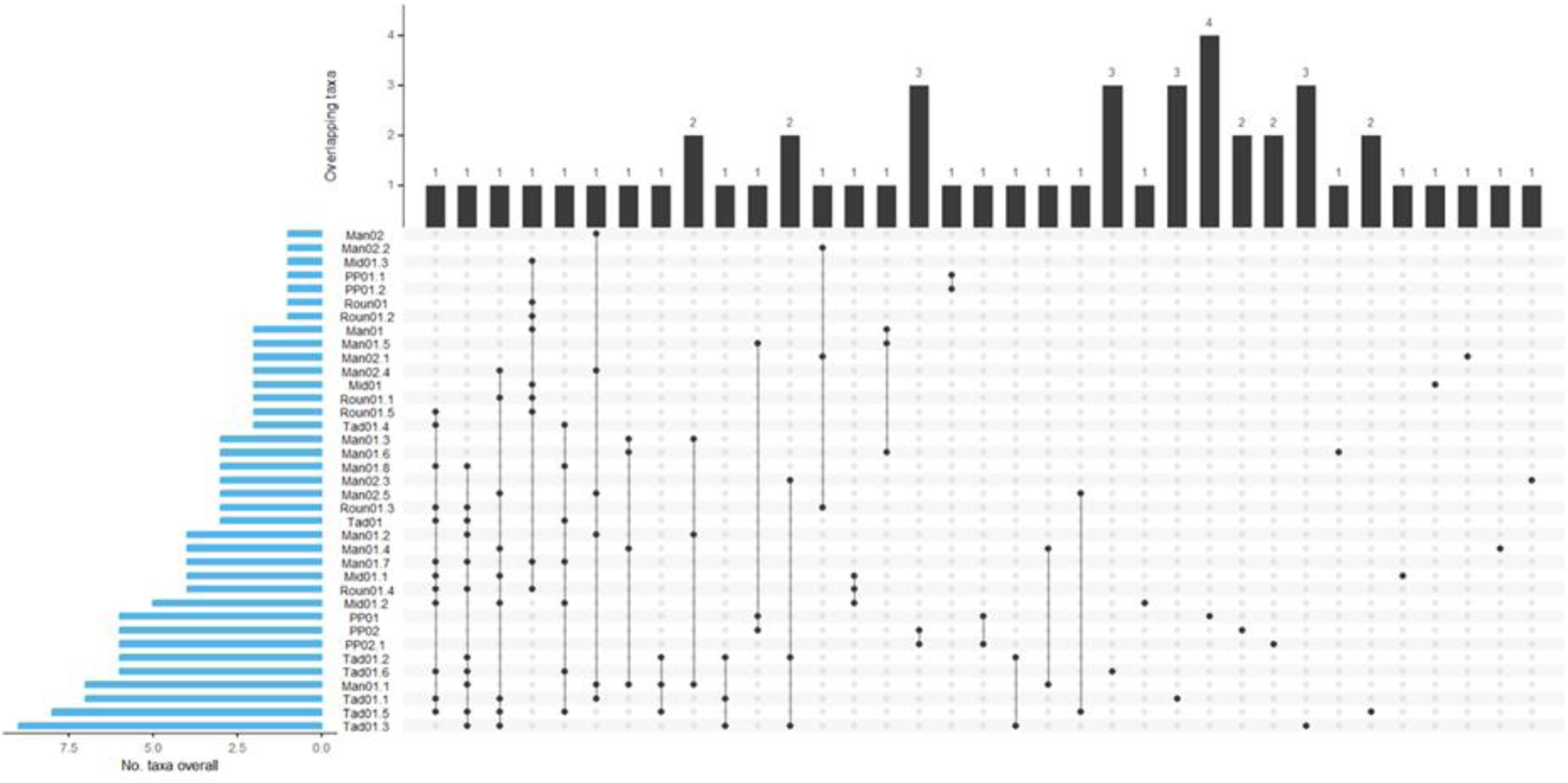
UpSet plots showing the number of ZOTUs shared between rock-holes for the uncollapsed *COI* eDNA metabarcoding data. Blue bars represent the total number of taxa within a single site. Black bars represent taxa that are shared between each site with black circles indicating which sites these taxa are shared with.

**Figure S5.**
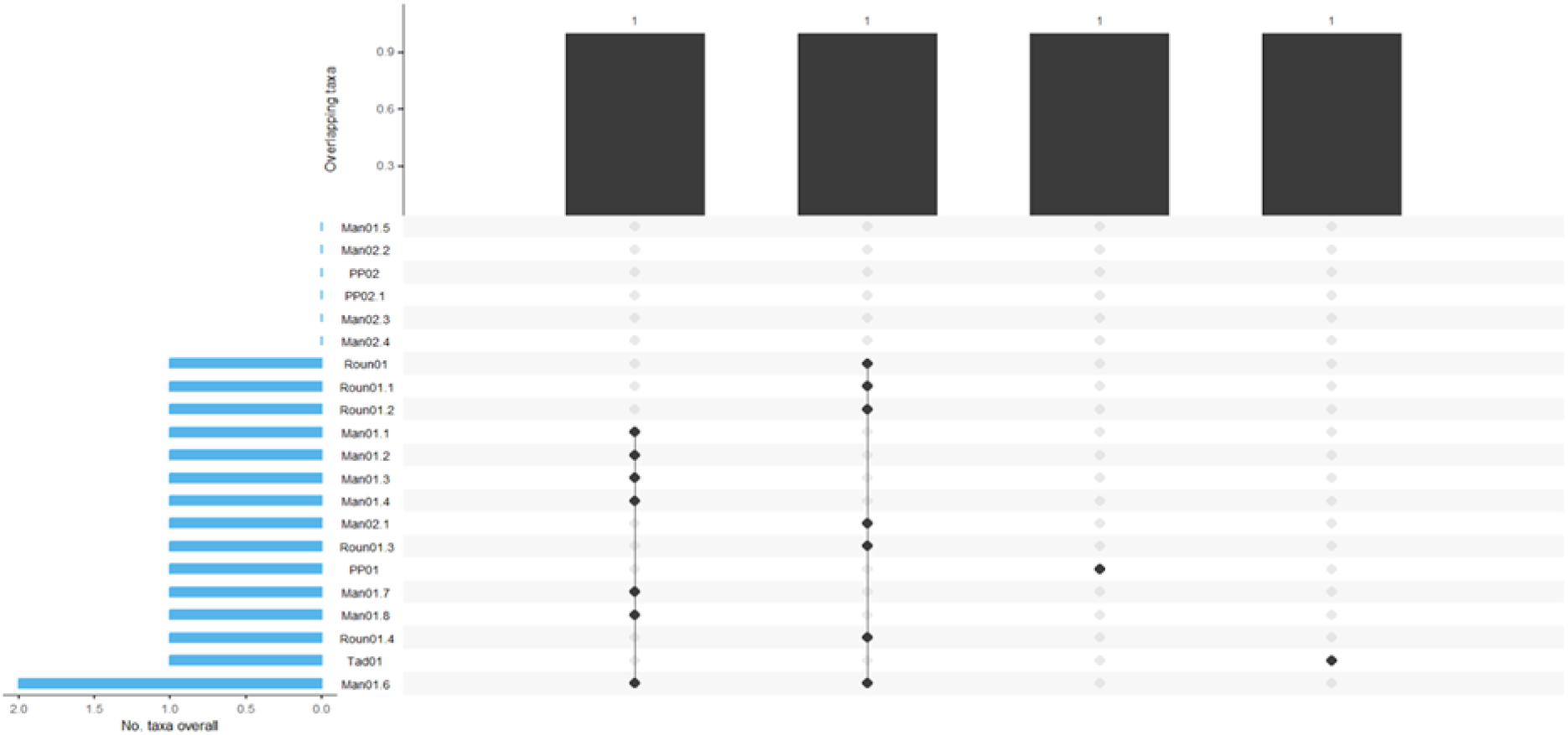
UpSet plots showing the number of ZOTUs shared between rock-holes for the uncollapsed *16S* eDNA metabarcoding data. Blue bars represent the total number of taxa within a single site. Black bars represent taxa that are shared between each site with black circles indicating which sites these taxa are shared with.

